# Discovery of molecular glues for modulating MHC presentation of cancer neo-antigens

**DOI:** 10.64898/2026.11.13.762685

**Authors:** Wenqi Shen, Ziyang Zhang, Changhao He, Keelan Z. Guiley, Vishal D. Lashkari, Julia Hernandez, Peter M. Bruno, Kevan M. Shokat

## Abstract

Antigen presentation by MHC class I molecules is essential for cancer immunosurveillance, but many cancer-specific antigens fail to form complexes with Class I MHC and are poorly presented on the cell surface. In this study, we present a chemical strategy to overcome this limitation by identifying peptide MHC glues (PMGs) that promote the formation of peptide-MHC complexes with otherwise unpresentable cancer antigens. Crystal structures of these complexes show that our compounds stabilize antigen-MHC binding by making simultaneous interactions with the MHC protein and the antigen peptides. These findings reveal a new approach for chemically rewiring Class I MHC antigen presentation and accessing rare cancer antigens for therapeutic targeting.

## Introduction

The presentation of intracellular peptides by Class I major histocompatibility complexes (MHC) is governed by the intrinsic affinity between a given peptide and the MHC allele ^1^. Among those peptides, neoantigens are peptides that contain somatic mutations specific to cancer cells. While other alterations such as splicing or post-translational modifications can also generate neoantigens, our study focuses on mutation-derived peptides. The presentation of neoantigens on the cell surface is important for recognition by cytotoxic T cells to achieve antitumor immunity ^2^. However, many neoantigens are not presented, often because they do not form stable complexes with MHC molecules ^3^. This substantially restricts the number of neoantigens that reach the cell surface ^4^.

Class I MHC molecules feature six conserved pockets (A–F) within the peptide-binding groove ^5^. High affinity binding of antigenic peptides to MHC-I is largely determined by the second (P2) and the last (PΩ) amino acids, known as “anchor residues”, which occupy the B- and F-pockets in the peptide binding groove, respectively. These two pockets are highly variable among the >10,000 polymorphic MHC alleles and shape the repertoire of antigens presented by each individual ^6^. For example, a mutant-specific antigen derived from KRAS G12V, (VVVGAVGVGK, aa7-16, hereafter referred to as V7-16), is a high-affinity binder to the HLA-A3 superfamily and has been targeted with cancer vaccines and protein-based therapeutics ^7^. However, the C-terminal lysine residue interacts poorly with the F-pocket of many common MHC alleles, limiting the utility of these therapies to a subset of patients ^8-10^.

To overcome this limitation, we envision that peptide-MHC interactions may be chemically stabilized with small molecule ligands to expand the landscape of cancer-specific antigens presented by MHC-I. Precedents for small molecules or metals which reshape the antigen specificity of MHC complexes have come from studies of drug hypersensitivity. For example, 5-8% patients treated with the anti-HIV drug abacavir developed hypersensitivity reactions ^11^. Although in the PREDICT-1 study, all confirmed cases of abacavir hypersensitivity occurred in HLA-B*57:01 carriers, only 47% of carriers developed the reaction, showing the HLA-B*57:01 allele is necessary but not sufficient for abacavir-induced hypersensitivity ^12,13^. The mechanism of this hypersensitivity was determined to be the binding of abacavir to HLA-B*57:01 and the “anchoring” of a self-peptide to the MHC molecule, which led to T cell recognition and hypersensitivity ^10,14^. Similarly, Clayton et al. characterized an “anchoring” example in their study of chronic beryllium disease ^15^. They found that patients with the HLA-DP2 allotype were at risk of developing hypersensitivity or autoimmunity when exposed to beryllium. Structural studies revealed that the Be^2+^ cation is buried in an acidic pocket formed between the DP2 β chain and the peptide antigen, templating an unnatural conformation change that triggers an immune response.

The inadvertent anchoring of peptides not normally presented by a particular HLA allele led us to ask if we could discover anchoring molecules for antigens of our choosing. We refer to these molecules as peptide MHC glues (PMGs). Here, we report the discovery of PMGs that enhance the presentation of potential cancer antigens derived from KRAS and p53 by HLA-B*57:01. We show that the PMG-MHC-peptide complexes are well-conformed and stable, and confirmed the PMG-induced cell surface presentation of KRAS and p53 neoantigens by mass spectrometry. Additionally, we solved X-ray co-crystal structures, further elucidating the molecular basis of molecular glue action. These studies provide a path for the discovery of PMGs for peptides of therapeutic interest.

## Results

Inspired by the molecular mechanism of abacavir-induced peptide presentation by HLA-B*57:01, we first asked whether abacavir can stabilize the binding of mutant KRAS antigens in this MHC allele, which has a valine at position 2 favored by the B-pocket and a lysine at the C-terminus, disfavored by the F-pocket. To quantitatively measure antigen binding, we adopted a sandwich ELISA that monitors the folding of MHC complexes with a conformation specific MHC-I antibody (W6/32) and a β2-microglobulin antibody (BBM.1) (Figure 1a) ^16,17^. We observed improved binding of a peptide with C-terminal valine (PepV, HSITYLLPV) in the presence of 100 µM abacavir, consistent with literature studies ^14^. However, abacavir failed to stabilize the mutant KRAS antigen V7-16 in this assay system, suggesting that the neomorphic surface created by abacavir in the HLA-B*57:01 F-pocket is not compatible with the mutant KRAS antigens (Figure 1b). We conclude abacavir is unlikely to be broadly suitable for the mutant cancer epitopes, though it may remain effective for epitopes with C-terminal isoleucine, leucine, and valine. Therefore, we set out to identify alternative molecular glues to anchor mutant KRAS antigens.

**Figure 1.**
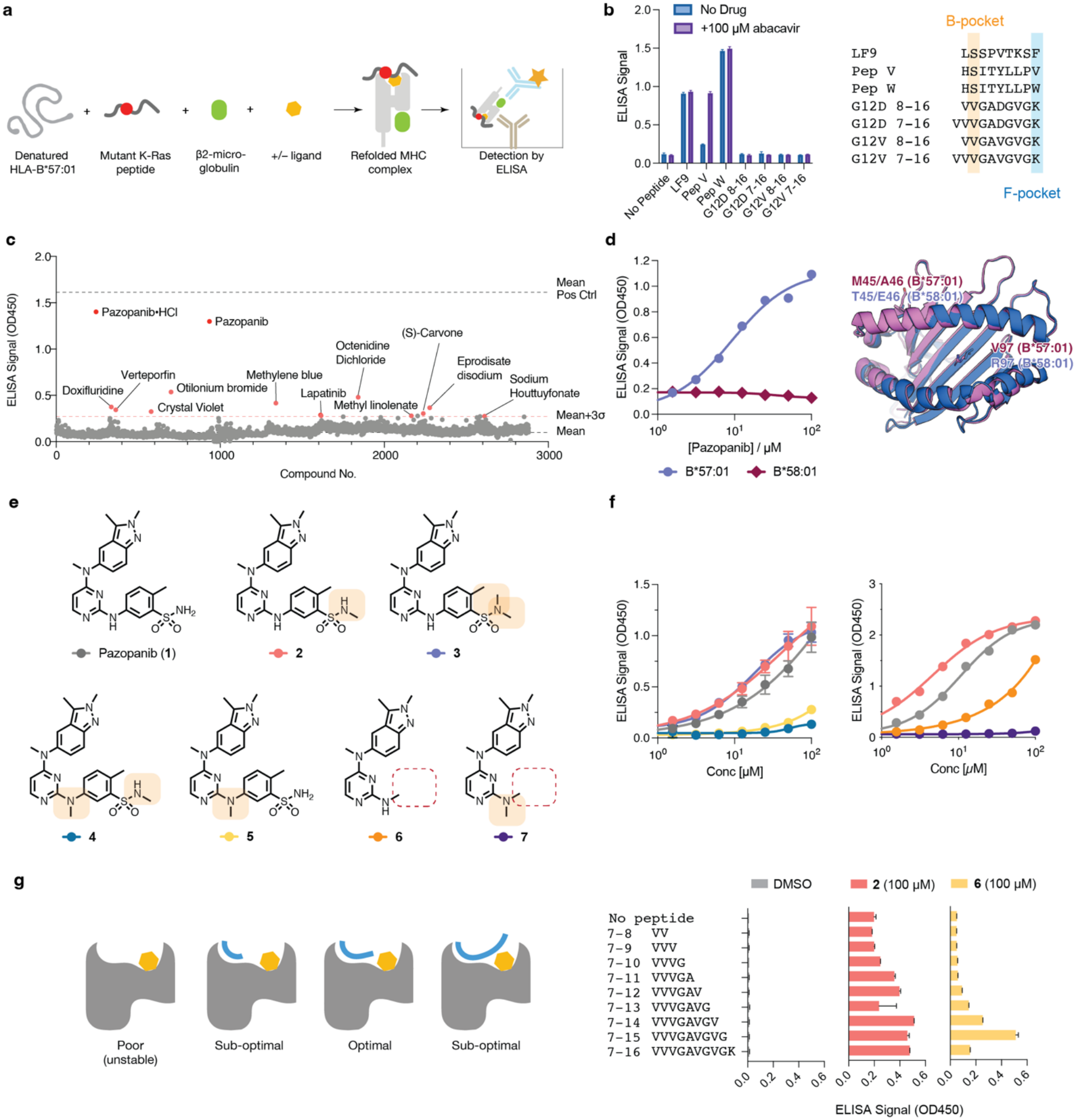
**a**. Schematic for detecting ligand-stabilized peptide binding to MHC-I with sandwich ELISA. **b**. Binding of peptide antigens to HLA-B*57:01 in the absence of presence of 100 µM abacavir. **c**. Chemical screen identifies pazopanib as a stabilizer for the binding of KRAS G12V 7-16 peptide to HLA-B*57:01. **d**. Pazopanib(**1**) stabilizes peptide binding to HLA-B*57:01 but not to HLA-B*58:01. **e**. Chemical structures of pazopanib and other synthetic analogs. **f**. Binding of KRAS G12V 7-16 peptide to HLA-B*57:01 in the presence of increasing concentrations of synthetic ligands. **g**. Cartoon model for compound-mediated stabilization of peptide-MHC complexes (left) and binding of the KRAS G12V 7-16 peptide of various lengths to HLA-B*57:01(right).

We hypothesized that structurally distinct small molecules may share a similar molecular mechanism as abacavir but be amenable to anchoring mutant KRAS peptides. To test this, we employed a refolding ELISA to screen an FDA-approved drug library (2880 compounds) using HLA-B*57:01 and the KRAS G12V peptide (VVVGAVGVGK). The screen led to the identification of one prominent hit – pazopanib (Figure 1c). Pazopanib was represented twice in the library (once as the free base and once as the HCl salt). While pazopanib promoted the refolding of HLA-B*57:01 in the presence of the KRAS peptide, it failed to do so with the closely related allele HLA-B*58:01 (Figure 1d). Because HLA-B*57:01 differs from HLA-B*58:01 at only three amino acids, of which only V97^B*57:01^/R97^B*58:01^ directly contact peptide antigens, we hypothesized that pazopanib specifically recognizes a pocket in HLA-B*57:01 near Val97. To further characterize the functional group(s) of pazopanib that contribute to the complex formation, we synthesized a small panel of six chemical derivatives of pazopanib and assessed their ability to stabilize the interaction between KRAS peptide and HLA-B*57:01 (Figure 1e-f). We first investigated the role of the sulfonamide group in pazopanib (**1**) by synthesizing its mono- and demethylated analogs **2** and **3**, respectively. While both compounds showed slightxly improved activity compared with pazopanib (**1**), we observed that the MHC complex formed with **2** was more stable: the complex remained intact after 72 h of incubation at 23ºC, whereas the complex formed with **1** was no longer detectable under the same conditions (SI Figure 1). We next asked whether the free N-H group attached to the 2-position of the pyrimidine ring was critical for its interaction with HLA-B*57:01. We prepared methylated analogs **4** and **5** and found that the both compounds had lost activity by ∼100-fold compared to their non-methylated counterparts (**4** vs **1** and **5** vs **2**). Such a drastic change suggests the loss of critical interaction mediated by the N-H group, on the order of ∼3 kcal/mol, which we hypothesized to be a hydrogen bond. Finally, removal of the entire 3-aminosulfonyl-4-methylphenyl group led to a decrease of activity, with compound **6** retaining measurable activity. Compound **7** which contains a dimethylamino group at the 2-position was inactive, consistent with the previous finding.

The F-pocket of HLA-B*57:01 favors bulky, hydrophobic residues such as tryptophan or tyrosine on the C-terminus of ligand peptides.^18^ However, our compounds overcome this preference, allowing a peptide with C-terminal lysine to bind. Together with the specific requirement for V97 in HLA-B*57:01 and the presence of the tryptophan-resembling dimethylindazole fragment in our synthetic ligands, our data led us to hypothesize that our compounds bind in the F pocket. To test this hypothesis, we assessed an array of progressively truncated versions of the V7-16 peptide to bind HLA-B*57:01 in the presence of our synthetic ligands (Figure 1g). Interestingly, compound **2** was able to stabilize most of the peptides in HLA-B*57:01 with a slight preference for the V7-14 8mer peptide, whereas the less sterically bulky compound **6** showed a clear preference for the longer V8-15 9mer peptide, supporting our proposed binding model (Figure 1g).

### Compound 2 binds in the F pocket of HLA-B*57:01

To understand the molecular basis of PMG binding with the KRAS G12V peptide and HLA-B*57:01, we sought to obtain a high-resolution structure HLA-B*57:01/compound **2**/KRAS peptide complex. Despite multiple attempts, we were unable to generate diffraction-quality crystals from this construct, likely due to its limited stability under crystal growth conditions. To overcome this limitation, we engineered an interchain disulfide bond between the α1 helix and the peptide antigen by introducing mutations A69C to the HLA-B*57:01 heavy chain and p6C to the peptide, respectively ^19^. The introduced disulfide bond increased the thermal stability of the ternary complex by 29.5ºC (SI Figure 2) and allowed us to solve a co-crystal structure at 1.7Å (Figure 2a). The overall architecture of the complex resembles typical Class I MHC complexes, with excellent electron density for the entire compound and the five N-term residues of the peptide clearly visible (Figure 2b).

**Figure 2.**
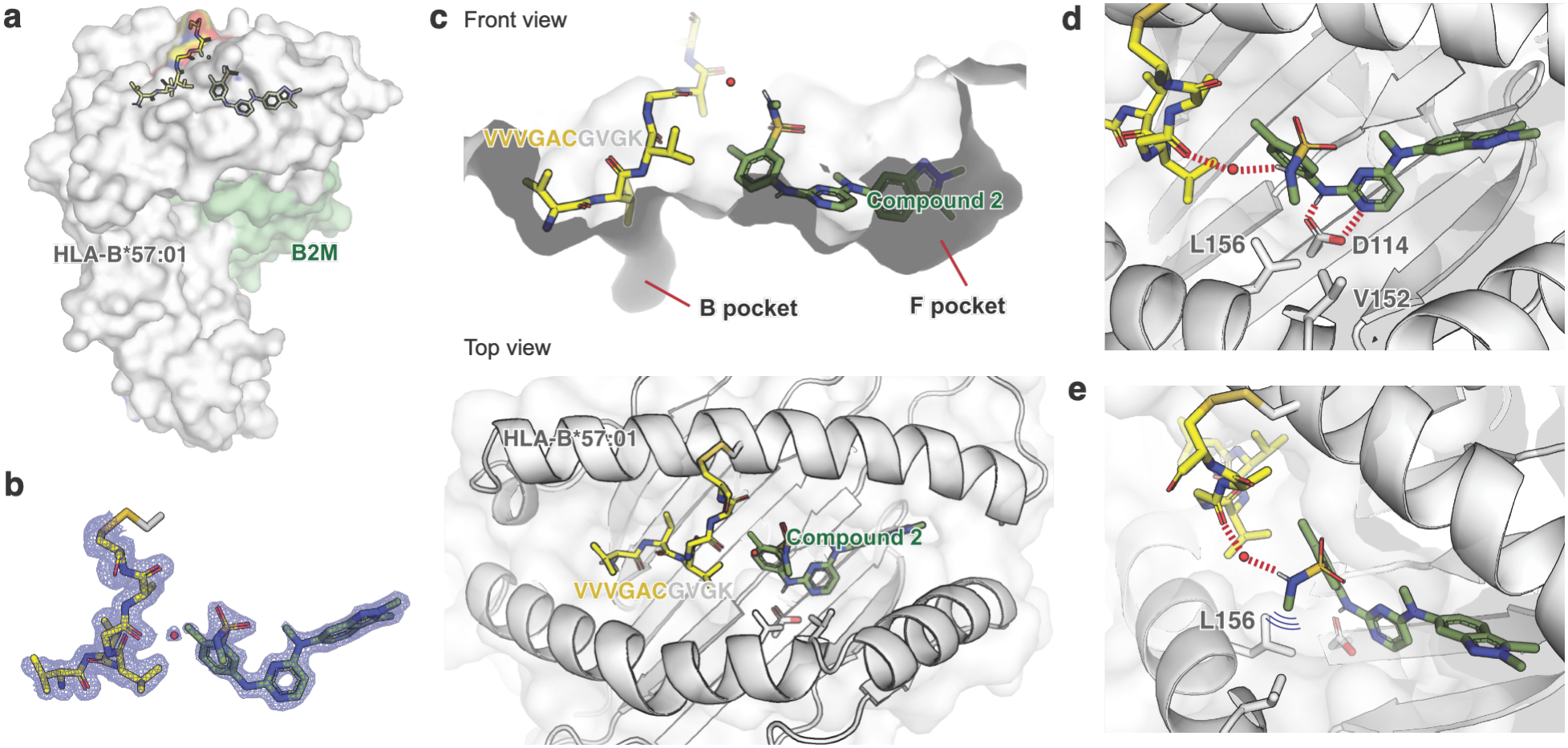
**a**. Overall architecture of the HLA-B*57:01•Compound 2•V7-16 complex. **b**. Fo-Fc map of the peptide and the chemical ligand, along with a key bridging water molecule. **c**. Front and top views showing the position of the peptide and compound **2. d-e**. Critical hydrogen bonding and hydrophobic interactions between the peptide, compound **2**, and the HLA-B*57:01 heavy chain.

Consistent with our hypothesis, compound **2** binds in the F-pocket, effectively mimicking the role of a terminal anchor residue and contributing to allele-specific binding. Compound **2** also simultaneously contacts the HLA-B*57:01 heavy chain and the KRAS peptide (Figure 2c). The bifurcate hydrogen bond between the aminopyrimidine of compound **2** and Asp114 of HLA-B*57:01 provides a mechanistic explanation of the importance of the free N-H group in our chemical exploration (Figure 2d). The indazole ring in compound **2** resides in a hydrophobic pocket, akin to how tryptophan side chains anchor natural antigens (Figure 3e) ^18^. We also observed a water-bridged interaction between the sulfonamide N-H of compound **2** and the amide backbone of the KRAS peptide (Figure 2d-e). Among all crystals we screened, we were unable to obtain resolvable density for the C-term portion of the peptide. This was not due to proteolytic cleavage of the peptide, as intact mass spectrometry analysis indicated that the peptide remained full-length (SI Figure 3). Instead, this likely suggests that the C-terminal residues interact weakly with the compound.

**Figure 3.**
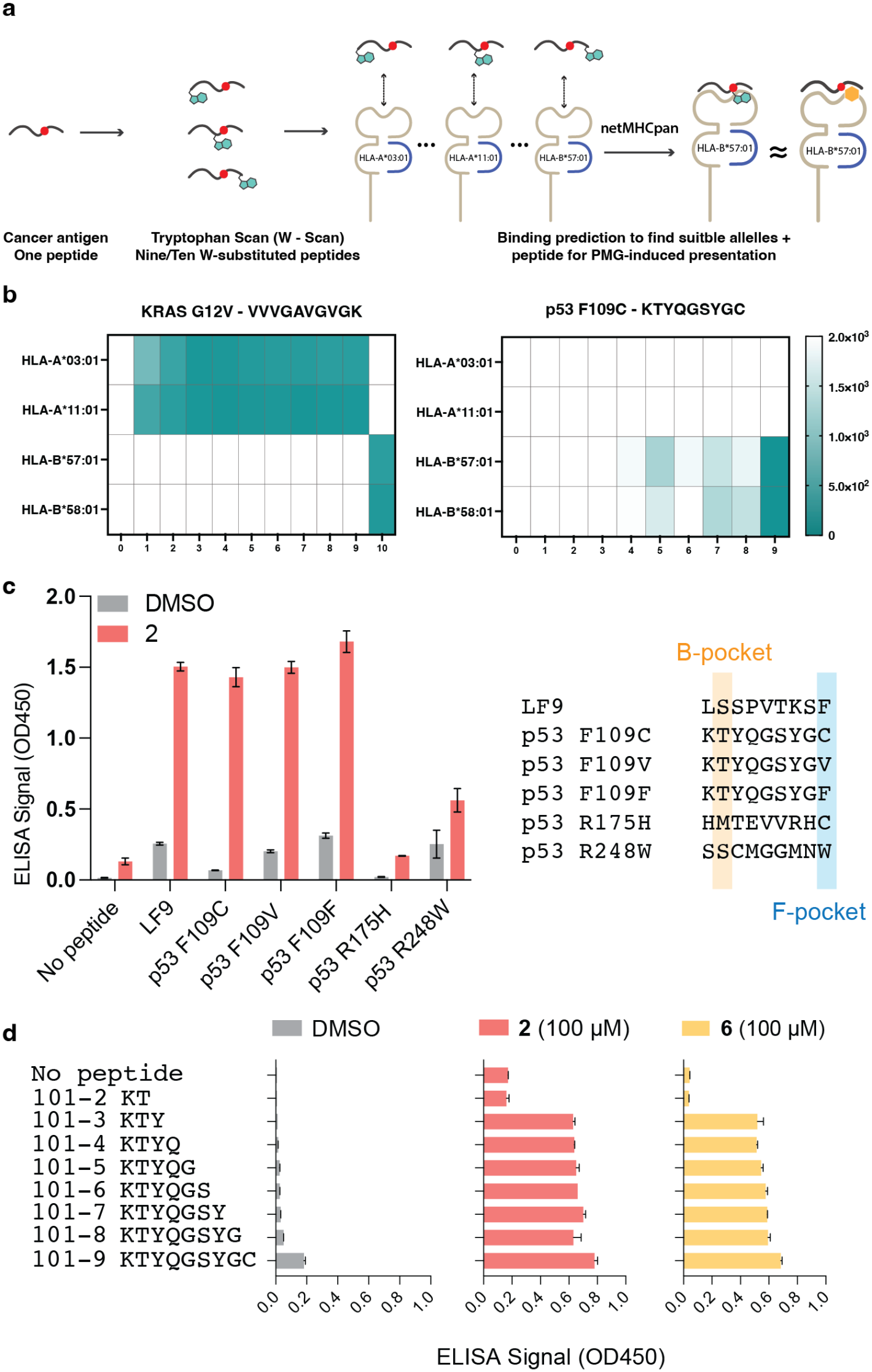
**a**. Schematic of W-scan. **b**. Heatmaps display NetMHCpan predicted binding affinities (in nM) for KRAS G12V (VVGAVGVGK) and p53 F109C (KTYQGSYGC) across four HLA class I alleles: HLA-A*03:01, HLA-A*11:01, HLA-B*57:01, and HLA-B*58:01. **c**. Binding of p53-derived peptide antigens to HLA-B*57:01 in the absence or presence of 100 µM compound **2**. Peptides tested include LF9 (LSSPVTKSF), p53 F109C (KTYQGSYGC), p53 F109V (KTYQGSYGV), p53 F109F (KTYQGSYGF), p53 R175H (HMTEVVRHC), and p53 R248W (SSCMGGMNW). **d**. Binding of p53 F109C-derived peptides of varying lengths to HLA-B*57:01.

### Stabilization of neoantigen from mutant p53

We next asked what other cancer neoantigens may be amenable to chemical stabilization in HLA-B*57:01. We chose to investigate p53 hotspot mutant neoantigens because p53 is the most frequently mutated gene in cancer, and due to the misregulation of MDM2, mutant p53 proteins are expressed at much higher level than the wildtype p53, providing an abundant source of antigenic peptides ^20^. With a large number of clinically observed p53 mutations, we sought to rank the “glueability” of mutant peptides *in silico* ^21^. Leveraging the neural-network-based epitope prediction algorithm NetMHCpan, we systematically substituted each residue of a mutant p53-derived with tryptophan, the bulkiest natural amino acid and preferred PΩ residue of HLA-B*57:01, and queried if HLA-B*57:01 could accommodate the resulting sequences (Figure 3a) ^22^. Our rationale was that a peptide MHC glue for mutant p53 antigens should phenotypically resemble a tryptophan substitution as we have observed in the crystal structure for the KRAS antigen. The small molecule would effectively act as the bulky tryptophan, allowing other residues to access the F-pocket. This approach was first used to assess abacavir self-peptide binding ^23^.

We included HLA-A*03:01 and HLA-A*11:01 based on prior reports that the KRAS G12V peptide binds these alleles ^24^. We also included HLA-B*58:01 alongside HLA-B*57:01 to test whether tryptophan substitution can differentiate binding preferences between these structurally related HLA-B molecules. We first performed this analysis on the KRAS V7-16 peptide and found that tryptophan substitution at P10 led to a predicted strong binder of HLA-B*57:01, consistent with our logic above. We then analyzed 3,183 patient-derived TP53 mutations from cBioPortal ^25-28^. After removing duplicates, we obtained 451 unique mutant protein sequences and generated overlapping 9-mer and 10-mer peptides encompassing mutation sites, resulting in 181 peptides per variant. Each peptide was analyzed by NetMHCpan to evaluate tryptophan-substituted variants for binding to HLA-B*57:01. We identified a subset of high-affinity binders, including p53 F109C (KTYQGSYGC), which corresponds to a known epitope region (KTYQGSYGF) documented in IEDB ^29,30^. The native F109C peptide showed weak predicted binding to HLA-B*57:01 and HLA-B*58:01, but strong binding was predicted upon tryptophan substitution at P9 (Figure 3b). We tested this experimentally using the refolding ELISA. While the p53 F109C peptide showed poor binding to HLA-B*57:01, compound **2** was able to improve its binding to a level similar to the positive control LF9. By contrast, two other peptides derived from common hotspot mutations, p53 R175H (167-175) and p53 R248W (240-248) were only marginally affected by compound **2** (Figure 3c). When we performed the peptide truncation scan for this F109C peptide, we observed a similar trend to that with K-RAS G12V peptide (Fig. 1g), where the signal is maximal with the 3mer peptide KTY (Figure 3d), suggesting a difference in the biochemical properties for this antigen.

We next obtained a 2.5-Å co-crystal structure of a complex formed by HLA-B*57:01, compound **2**, and p53 F109C(101-109) peptide without the need to introduce an artificial disulfide bond required in the case of the K-Ras (G12C) peptide. The overall architecture resembles the complex with KRAS G12C(7-16), and compound **2** adopts an identical conformation as seen in the earlier structure (Figure 4a). In the non-covalent complex, the p53 peptide was also only partially resolved up to residue P5, and its backbone amide is positioned closer to the peptide binding groove (Figure 4b), likely due to the lack of forced disulfide formation in the KRAS complex. We compared the conformation of the antigenic peptides in our two structures with that of a natural antigen (PDB: 2RFX) and an abacavir-induced self-antigen (PDB: 3UPR) (Figure 4c). While residues P1-P3 were almost identically positioned in these four structures, we observed prominent differences starting from P4. Specifically, abacavir led to rearrangement of the P5-P6 region but allowed the C-terminus of the peptide to bind, whereas compound **2** completely displaced the C-terminal portion of peptide backbone. Such a difference is likely the result of the bulky indazole group but suggests the possibility that replacing the indazole ring with smaller substituents may allow the accommodation of the C-terminal residues, possibly with sequence specificity.

**Figure 4.**
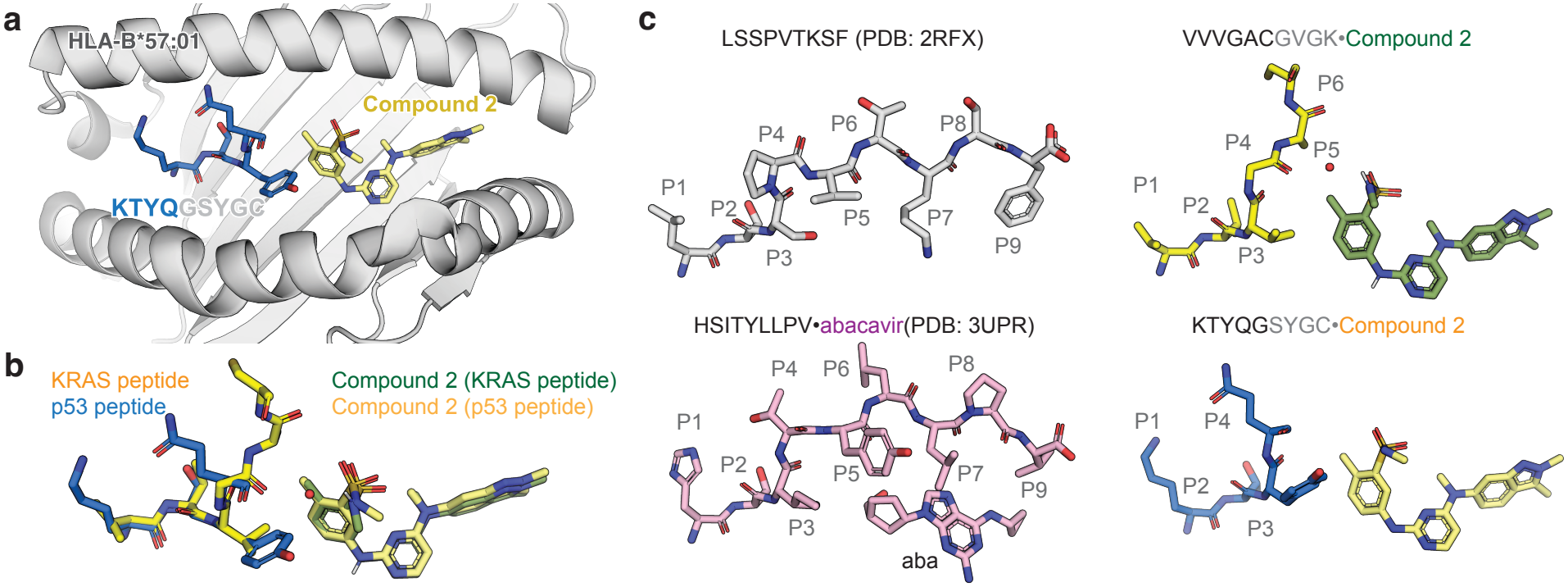
**a**. Co-crystal structure of HLA-B*57:01 with compound **2** and p53 F109C(101-109) peptide. **b**. Overlaid peptide and compound conformations of the complexes formed with mutant KRAS and p53 peptides. For clarity, HLA-B*57:01 protein was omitted from the plot. **c**. Comparisons the peptide conformations in HLA-B*57:01 complexes of a natural antigen (PDB:2RFX), an abacavir-induced self-antigen (PDB: 3UPR), and compound **2**-stabilized mutant KRAS and p53 antigens (this study).

Our structure revealed both similarities and differences to abacavir-bound HLA-B*57:01: while both compounds bind in the F-pocket of HLA-B*57:01, abacavir does not hydrogen bond with Asp114, but its cyclopropyl group makes hydrophobic contacts with the Leu/Val side chain of the antigen peptide, potentially explaining its inability to anchor the KRAS peptides which contains a C-terminal lysine residue.

### PMGs increase the presentation of neoantigens on the cell surface

We next sought to assess whether compound **2** facilitates the presentation of mutant KRAS and p53 antigens on the cell surface using targeted mass spectrometry.^31^ We treated COS-7 cells ectopically expressing HLA-B*57:01 and KRAS G12V (Figure 5a) with DMSO or compound **2** (10 µM), enriched MHC-bound peptide by immunoprecipitation with the pan-HLA antibody W6/32, and monitored the levels of KRAS G12V(7-16) peptide by tandem-mass spectrometry. Since COS-7 cells do not express human MHC alleles, this method allows the allele-selective analysis of surface antigen levels. We identified the G12V(7-16) peptide with a full series of y ions (Figure 5b) and observed treatment-dependent increase of three representative daughter ions (Figure 5c). We next repeated the targeted mass spectrometry experiment by ectopically expressing HLA-B*57:01 and p53 F109C (Figure 5d). We were able to identify the F109C peptide (Figure 5e) and observed a similar treatment-dependent increase pattern of three representative daughter ions (Figure 5f).

**Figure 5.**
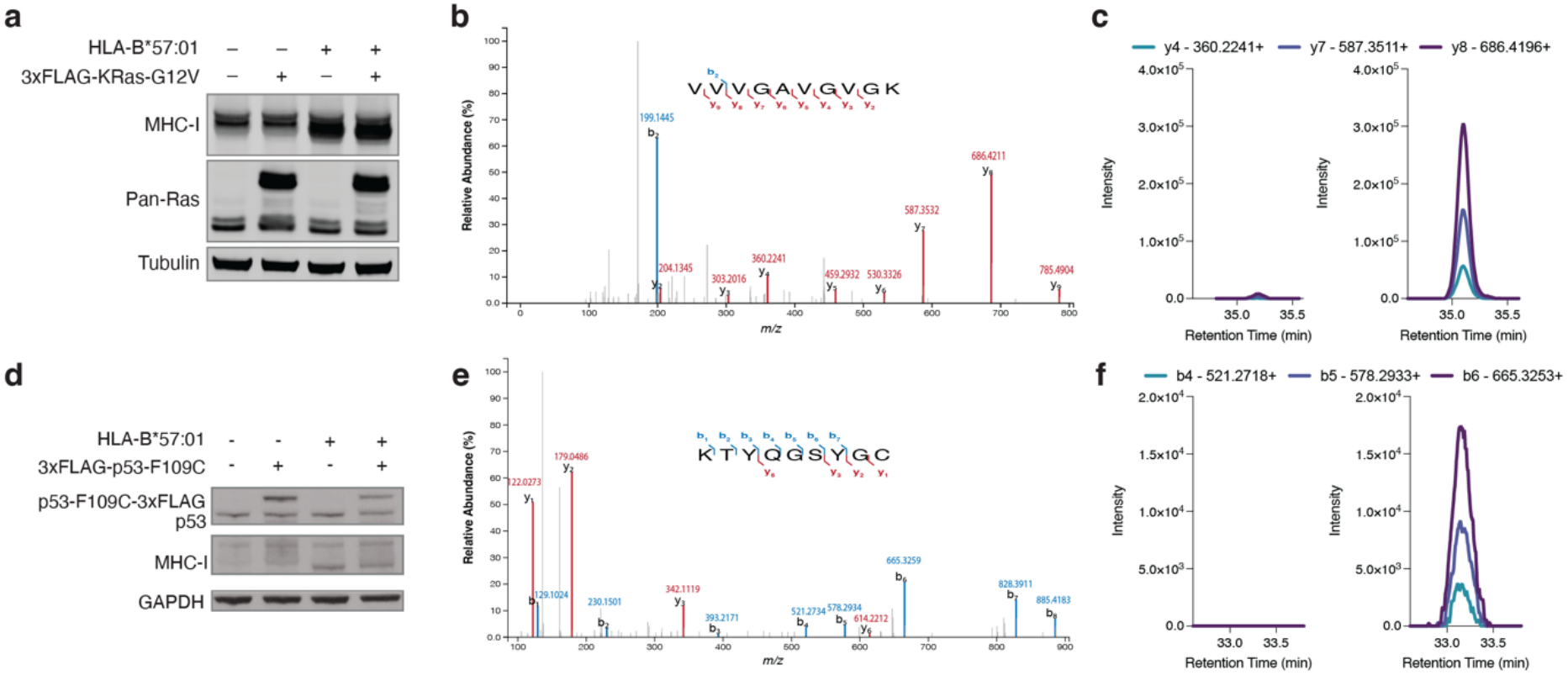
**a**. Expression of HLA-B*57:01 and K-Ras G12V in COS-7 cells. **b**. MS2 spectrum of KRAS G12V(7-16) peptide with key b and y ions highlighted. **c**. Intensities of three representative MS2 daughter ions. **d**. Expression of HLA-B*57:01 and p53 F109C in COS-7 cell. **e**. MS2 spectrum of p53 F109C (101-109) with key b and y ions highlighted. **f**. Intensities of three representative MS2 daughter ions quantified by parallel reaction monitoring of the parent peptide ion and MS2 extraction.

### Immunopeptidomics reveals minimal compound-induced global changes to the peptide repertoire

We asked whether and how compound **2** reshapes the repertoire of HLA-B*57:01 antigens presented on the cell surface beyond the mutant KRAS and p53 epitopes. To profile the compound-induced changes in the immunopeptidome, we first employed the EpiScan platform, a functional genetic, quantitative method for high-throughput MHC-I ligand discovery ^32^. EpiScan cells are derived from HEK293T but lack TAP1/2, ERAP1/2, and endogenous MHC-I, resulting in reduced surface MHC levels even after re-introduction of a single allele of interest. Exogenous peptides expressed via lentiviral transduction will restore surface levels only if the peptide binds to the re-introduced allele, and such peptides can be identified at scale via next-generation sequencing. We used EpiScan cells expressing only HLA-B*57:01 and then transduced them with a library (170,000 sequences) containing peptides enriched for binding to HLA-B*57:01, thus biased toward preferred A-, B-, and F-pocket residues, but relatively randomized at the remaining positions. Cells were treated with DMSO, abacavir, or compound **2**, sorted for restored surface MHC levels, and processed for next-generation sequencing to identify the peptides that bound to HLA-B*57:01 under each treatment condition (Figure 6a). Compound **2** treatment led to a global increase in MHC expression in GFP^+^ cells (cells expressing a library peptide) as well as GFP– cells (unselected cells not transduced with a library peptide) (Figure 6b). To validate this observed population shift, we treated the parental EpiScan cells containing HLA-B*57:01 (no peptide library transduction) with the same conditions and quantified surface MHC levels (Figure 6c). We again saw a consistent increase in surface MHC levels after treatment with compound **2**, as was observed in the library screen and our previous ELISA experiment, confirming the role of compound **2** in the MHC shift.

**Figure 6.**
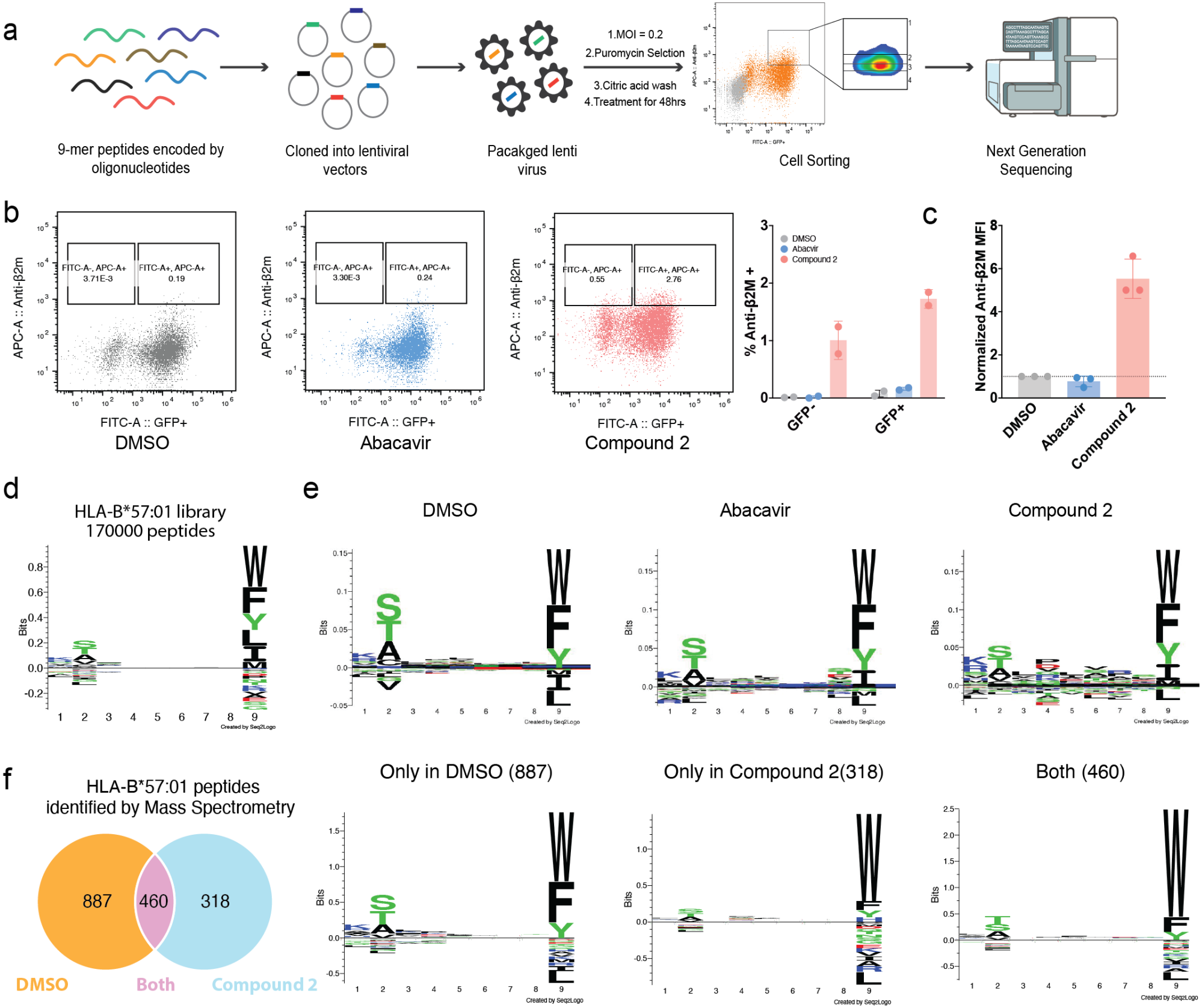
EpiScan reveals global changes in surface peptides upon compound 2 treatment. **a**. EpiScan workflow. **b**. Flow cytometry and quantification of surface MHC levels of cells used in EpiScan screen following drug treatment. The black population represents untreated non-transduced input cells that do not contain the peptide library. Colored populations represent cells treated with DMSO (grey), 10 μM Abacavir (blue), or 10 μM Compound **2** (coral). **c**. Quantification of surface MHC levels in the non-transduced parental cells treated with DMSO, 10 μM abacavir, or 10 μM compound **2. d**. Logo plot showing the composition of the peptide library with 170,000 peptides. **e**. Logo pots of peptides identified in each condition with corrections applied based on the position-specific frequencies of the library. **f**. Untargeted immunopeptidome isolated from 721.221-HLA-B*57:01 cells and analyzed by mass spectrometry. Venn diagram showing the number of unique HLA-B*57:01-associated peptides identified by untargeted in DMSO (orange, 887 peptides), compound 2–treated cells (blue, 318 peptides), or shared between both conditions (overlap, 460 peptides). Sequence logos show the position-specific amino acid composition of peptides identified only in DMSO, only in compound **2**, or in both conditions.

To understand the molecular basis of this restoration, we performed genomic sequencing to identify peptides enriched in cells exhibiting restored surface MHC levels. After sequencing the genomic DNA of the cells with restored surface MHC levels, peptide binders to HLA-B*57:01 were identified. To identify preferred residues at each position, we determined peptide preference relative to the input library (Figure 6d) to generate logo plots for each treatment condition (Figure 6e). As expected, we observed a similar, dominant peptide presentation pattern across all three conditions, reflecting the untreated HLA-B*57:01 peptide preference. However, compound **2**-treated EpiScan screens revealed two major trends: 1) the dominant peptide preference was similar to that of the untreated screen, reflecting a more accommodating F pocket, and 2) Compound **2** did not exert a consistent effect on allele-specific binding preferences. The increased resemblance of the peptides to the input library indicates a potential nonspecific effect on presentation.

In parallel, we performed untargeted immunopeptidome analysis by eluting peptides from immunoprecipitated MHC and then analyzing them via tandem mass spectrometry. This was undertaken on a monoallelic stable cell line expressing HLA-B*57:01 derived from 721.221 cells, which are HLA-A/B/C-null but contain an intact antigen processing machinery, and treated with DMSO or compound **2** (Figure 6f) ^31,33^. We found that the peptide repertoire under compound 2 treatment overlapped with the DMSO control by 460 peptides. We also observed condition-specific peptides, with 318 peptides unique to the Compound 2 treated samples and 880 peptides unique to DMSO. Despite the identification of 318 compound 2 specific peptides, their position specific amino acid preferences remained consistent with the canonical HLA-B*57:01 motif.

Together with our biochemical assay data, these findings show that compound **2** allows presentation of additional peptides by HLA-B*57:01 without compromising its ability to present native antigens.

### Structure-guided optimization decouples MHC binding and kinase inhibition

One limitation of our compounds was that they derive from the pan-tyrosine kinase inhibitor pazopanib, which may inhibit kinase-mediated signal transduction critical for T cell function at the doses required for MHC binding. Our attempt to modify the hinge-binding motif (compounds **4** and **5**) to diminish kinase inhibition,however, led to a loss of HLA-B*57:01 binding. A comparison of our crystal structures with the abacavir led to the observation that the NH_2_ group in abacavir forms a hydrogen bond with the backbone amide of Ile124, an interaction does not present with compound **2**. We hypothesized grafting an NH_2_ group at the equivalent position on compound **2** might improve its potency as an MHC anchor but interfere with kinase binding (Figure 7a). To test our hypothesis, we synthesized compound **8** and found that it maintained the ability to stabilize the mutant KRAS peptide in HLA-B*57:01 but showed substantially decreased activity in inhibiting the tyrosine kinases we surveyed (Figure 7c). Whereases, compound **2** inhibits several T cell kinases such as Lck and Fyn at sub-micromolar concentrations in biochemical kinase assays (Figure 7c).To test whether compound **8** suppresses the activation of primary T cells, we stimulated PBMCs from an HLA-B*57:01 positive donor with anti-CD3 and anti-CD28 antibodies and measured several cell surface markers of activation. At 10 µM concentration, compound **2** inhibited T cell activation (quantified by the frequency of CD137+ CD69+ CD8 T cells) by >90%, whereas compound **2** showed <50% inhibition. At a 1 µM concentration, T cell activation was not affected by either compound (Figure 7d). While compound **8** does not show perfect selectivity, our data show that it is possible to decouple kinase inhibition from MHC anchoring activity through structure-informed chemical optimization.

**Figure 7.**
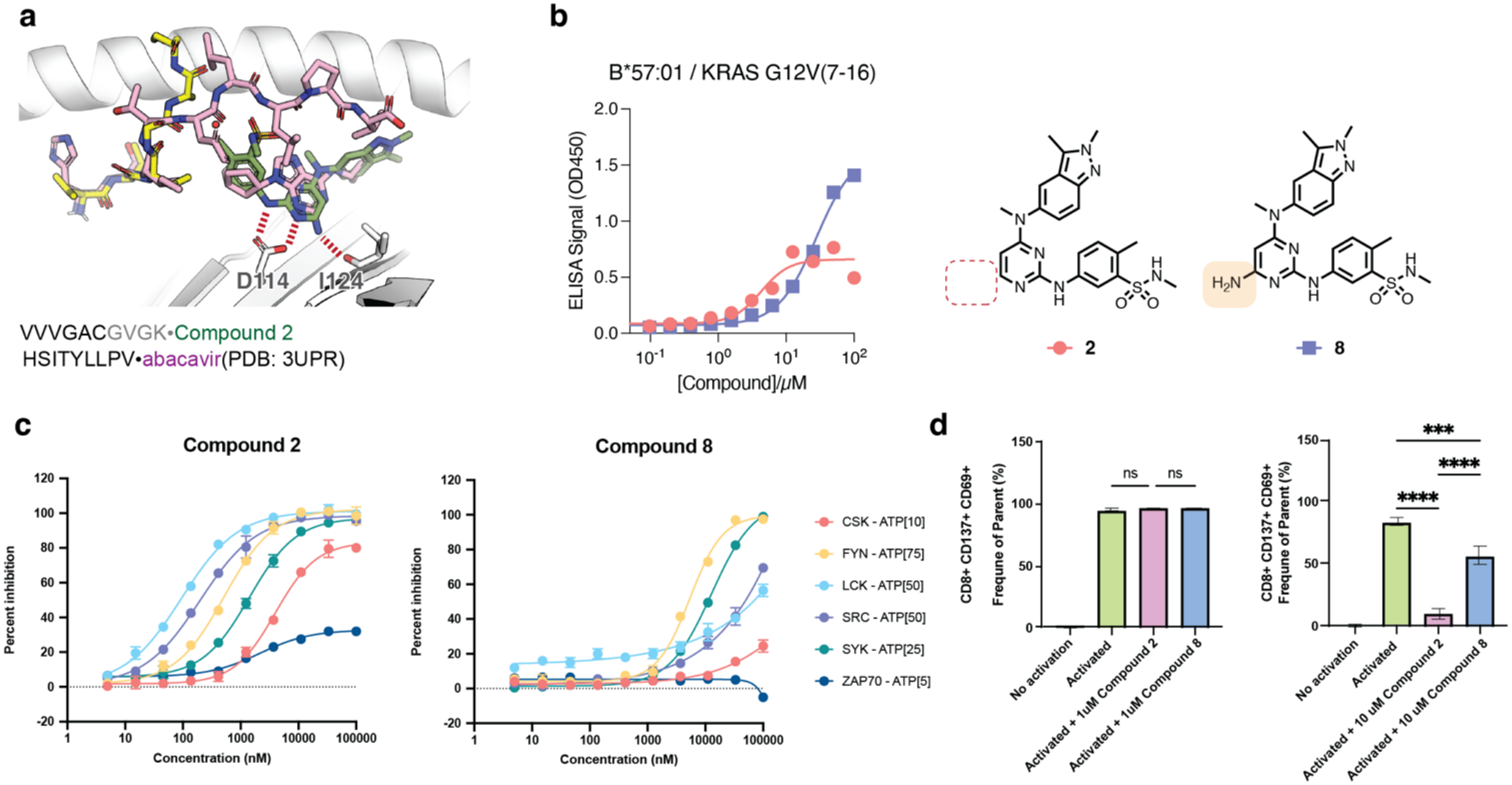
Design of PMGs with reduced inhibition of Src family kinases and reduced T cell suppression. **a.**Overlaid PDB 3UPR with the structure of HLA-B*57:01 with compound **2** and mutant KRAS peptide **b**. Refolding ELISA data with the new analogs. **c**. Inhibition of select Src-family kinases involved in T cell signaling by compounds **2** and **8. d**. T cell activation assay performed using human peripheral blood mononuclear cells (PBMCs) stimulated with anti-CD3 and anti-CD28 antibodies in the presence of 1 µM or 10 µM of compound 2 or compound 8.

## Conclusion and Discussion

In this work, we report the discovery and structural characterization of novel molecular glues that modulate Class I MHC peptide presentation. These compounds highlight the potential of small molecules to reshape the repertoire of MHC antigens through neomorphic interactions. By stabilizing poorly presented cancer neoantigens in an MHC-restricted fashion, these compounds hold promise as pharmacological agents to enhance tumor immune recognition. We know from cases of exceptional checkpoint blockade responders which harbor rare driver oncogene mutations such as BRAF (N581I) or AKT (E17K), deep long term responses in these patients were due to T cells specifically able to detect peptides presented by class I MHC molecules ^34^. These patients’ tumors lacked high mutational burdens which suggests that eliciting responses to oncogenic driver neoantigens could greatly expand the utility of current checkpoint therapies.

EpiScan provides a scalable, functional readout of peptide binding to a single HLA allele, but it has limitations that can influence interpretation. Because the system was designed to only exclude endogenous peptides longer than eight amino acids, shorter host-derived peptides may still load onto HLA-B*57:01 and may be preferentially stabilized by compound **2**, potentially contributing to the apparent enrichment profile and the observed increase in surface MHC even in the absence of library expression(Fig. 6b,c). In addition, the W6/32-based detection reports β2m–associated heavy-chain assembly rather than the peptide-binding region, so it cannot distinguish complexes containing peptide alone from those containing peptide plus compound or complexes stabilized predominantly by compound. These assay-level features motivated the use of complementary approaches to define how compound **2** influences peptide loading and complex stability. Notably, untargeted MHC immunoprecipitation coupled to mass spectrometry in monoallelic 721.221-HLA-B*57:01 cells independently showed that compound **2** largely maintained the canonical HLA-B*57:01 motif, reaching the same overall conclusions as EpiScan. Together, these orthogonal datasets indicate that compound **2** broadly boosts HLA-B*57:01 presentation without eroding allele-specific preferences, while highlighting the value of integrating functional and biochemical measurements.

Our data also suggests a possible explanation for the clinically observed liver toxicity associated with Pazopanib^35^. Pazopanib may increase the presentation of certain liver-specific antigens in an immunocompetent conformation, triggering T cell response.

Further investigation into the mechanisms underlying peptidome changes will be crucial for refining PMG design for therapeutic applications.

## Notes

### Competing Interest Statement

K.M.S. and Z.Z are inventors on patents owned by University of California, San Francisco covering stabilization of MHC complexes. P.M.B. is co-inventor of EpiScan PCT/US2020/062912. P.M.B. also serves on the scientific advisory board of and holds equity in Sift Biosciences. Z.Z. has consulting agreements for the following companies, which involve monetary and/or stock compensation: X-biotix, Montara Therapeutics, Kimia Therapeutics, Prelon Therapeutics. K.M.S. has consulting agreements for the following companies, which involve monetary and/or stock compensation: BridGene Biosciences, Erasca, Exai, G Protein Therapeutics, Genentech, Kumquat Biosciences, Kura Oncology, Lyterian, Merck, Montara Therapeutics, Nextech, Revolution Medicines, Rezo, Tahoe, Totus, Type6 Therapeutics, Wellspring Biosciences (Araxes Pharma).

### Funding

- Howard Hughes Medical Institute
- The Mark Foundation for Cancer Research
- Damon Runyon Cancer Research Foundation: DRG-2281-17, DRG-2399-20
- Life Sciences Research Foundation
- National Cancer Institute: P30CA082103
- National Institutes of Health: 1S10OD028511-01

